# Neural network models for spinal implementation of muscle synergies

**DOI:** 10.1101/2021.10.27.466061

**Authors:** Yunqing Song, Masaya Hirashima, Tomohiko Takei

**Affiliations:** Graduate School of Medicine, Kyoto University; Center for Information and Neural Networks (CiNet), Advanced ICT Research Institute, National Institute of Information and Communications Technology (NICT); Graduate School of Frontier Biosciences, Osaka University; The Hakubi Center for Advanced Research, Kyoto University; Brain Science Institute, Tamagawa University

**Keywords:** muscle synergies, optimization, spinal interneurons, neural manifold, redundancy

## Abstract

Muscle synergies have been proposed as functional modules to simplify the complexity of body motor control; however, their neural implementation is still unclear. Converging evidence suggests that output projections of the spinal premotor interneurons (PreM-INs) underlie the formation of muscle synergies, but they exhibit a substantial variation across neurons and exclude standard models assuming a small number of unitary “modules” in the spinal cord. Here we compared neural network models for muscle synergies to seek a biologically plausible model that reconciles previous clinical and electrophysiological findings. We examined three neural network models: one with random connections (non-synergy model), one with a small number of spinal synergies (simple synergy model), and one with a large number of spinal neurons representing muscle synergies with a certain variation (population synergy model). We found that the simple and population synergy models emulate the robustness of muscle synergies against cortical stroke observed in human stroke patients. Furthermore, the size of the spinal variation of the population synergy matched well with the variation in spinal PreM-INs recorded in monkeys. These results suggest that a spinal population with moderate variation is a biologically plausible model for the neural implementation of muscle synergies.

## 1. Introduction

Our body is remarkably complex, yet we display a highly stable motor performance. For example, to reach for a coffee cup on a desk, there are an infinite number of patterns of muscle activity involved in extending the arm because multiple muscles span the same joint. Nevertheless, we show a highly stereotyped movement trajectory and agonist-antagonist muscle activity patterns^1^. Understanding how the central nervous system (CNS) coordinates the redundant musculoskeletal system is a central question of motor control.

Muscle synergies have been proposed as a solution to control redundant systems by coordinating a number of muscles with a smaller number of control modules, which is called muscle synergies^2-4^. This hypothesis is phenomenologically supported by experimental observations that a linear combination of basic patterns of muscle activity successfully reconstructs muscle activity during a wide range of behaviors, including reflex movements^2,5^, postural tasks^6-8^, locomotion^9,10^, reaching, and grasping^11-13^. However, there is still a heated debate regarding whether these experimental observations reflect a physiological basis of low-dimensional control in the CNS or an epiphenomenon based on task constraints and/or biomechanics^14-17^.

Hirashima and Oya (2016) demonstrated that a neural network model that did not explicitly assume muscle synergies in the model could produce a synergy-like low-dimensional structure in muscle activity when the network was optimized to produce different combinations of elbow and shoulder torques while minimizing motor effort and motor error^16^. This clearly shows that detecting muscle synergies in muscle activity does not itself provide evidence for the existence of basic modules in the control system. Therefore, to examine the existence of muscle synergies, it is essential to identify the neural implementation of muscle synergies in the nervous system and develop a biologically plausible model.

Accumulating evidence from physiological and anatomical studies suggests that output projections of spinal premotor interneurons (PreM-INs) to motoneuron pools are the neural basis of muscle synergies in frogs^18^, rodents^19^, and primates^13,20,21^. Our previous study demonstrated that spinal PreM-INs have a divergent projection to multiple hand and arm motoneurons^20,21^ and their spatial distribution corresponds to the spatial weight of muscle synergies extracted from muscle activity. These results suggest the contribution of spinal PreM-INs to the generation of muscle synergies^13^. However, while the output projection of each PreM-IN corresponded to the muscle synergies, there was a substantial variation in the projection patterns across PreM-INs. This finding clearly contradicts the assumption that each muscle synergy can be modeled as a unitary “module,” which is often used in muscle synergy models. Therefore, the development of neural models to explain how muscle synergies are implemented with divergent spinal PreM-INs has been awaited.

Here, we created neural network models for muscle synergies and compared their performance to explain a known experimental phenomenon: the robustness of muscle synergies for cortical stroke^22,23^. In experiment 1, we examined the existence of muscle synergies by comparing two neural network models: one with random connections from the cortical layer to the muscle layer (non-synergy model) and the other with a small number of spinal synergies in the middle (simple synergy model). Then, in experiment 2, we sought a more biologically plausible model for muscle synergies by allowing a certain level of variation in spinal neurons that constitute muscle synergies (population synergy model). Our results demonstrate that 1) the simple and population synergy models emulate the robustness against cortical stroke that was observed in human stroke patients, and that 2) the population synergy model has a similar performance to the simple synergy model when the spinal variation was moderate, as in the variations of spinal PreM-IN recorded in non-human primates in our previous study. These results suggest that the population synergy model is a biologically plausible model for neural implementation of muscle synergy to achieve robust motor control.

## 2. Methods

### 2.1 Neural network models

We compared three types of neural network models for neural implementation of muscle synergy:1) non-synergy model, 2) simple synergy model, and 3) population synergy model.

#### 2.1.1 Non-synergy model

The non-synergy model was the same as the neural network model in previous studies^16,24^. Briefly, a linear neural network was used to convert the desired torque (input layer) into the actual torque (output layer) through an intermediate layer composed of 1,000 cortical neurons and eight muscles (Fig. 1a). Each cortical neuron received the desired torque vector (*τ*) from the input layer with synaptic weights (*W*). Then, the cortical neurons projected to the muscles with innervation weights (*Z*). The innervation weights from each cortical neuron to muscles were established so that *Z*_*i*_ (*i* = 1–1,000) were uniformly distributed on the surface of a sphere in an 8-dimensional space, with a radius of 0.002 (= 2/*n*). The eight muscles included two shoulder flexors (outer and inner shoulder flexors: SFo and SFi, respectively), two shoulder extensors (outer and inner shoulder extensors: SEo and SEi, respectively), an elbow flexor and extensor (EF and EE, respectively), and a biarticular flexor and extensor (BiF and BiE, respectively). The mechanical effects of the muscles (mechanical direction vectors, MD vectors) were in line with those of the previous model (*M*)^16,24^. Muscle activity (*a*) was constrained to be positive, thereby making the model non-linear. The total output of the network (*T*) was expressed as the vector sum of the mechanical output of all muscles. The synaptic weight from the input layer to the cortical neurons was initially selected from a standard normal distribution with a zero mean and unitary standard deviation. Then, it was modified by the error feedback-with-decay algorithm^16,24^ (see Training procedure).

**Fig. 1.**
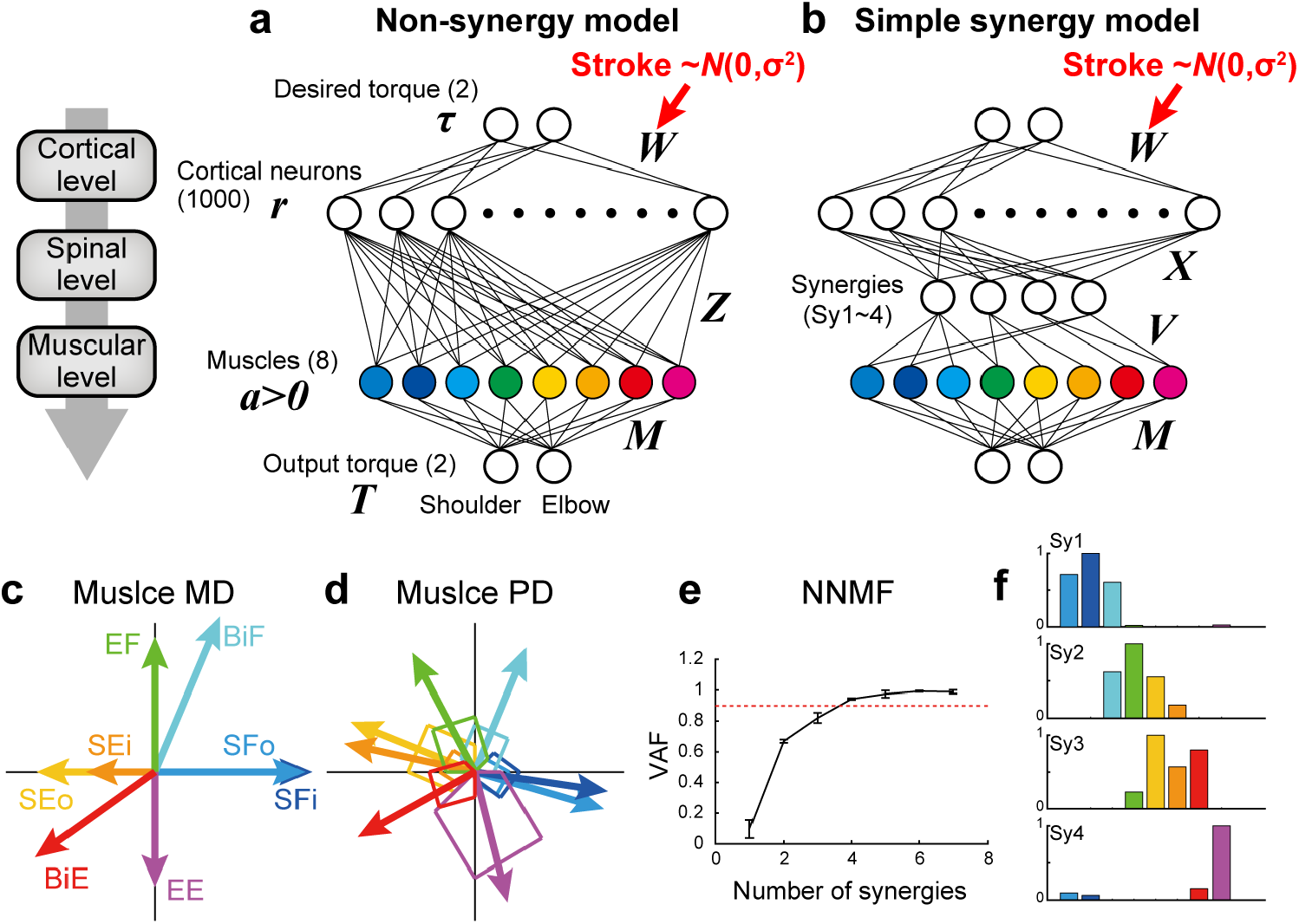
Non-synergy and simple synergy models for shoulder and elbow torque task. **a-b**. Neural network model without (non-synergy model, **a**) or with a synergy layer (simple synergy model, **b**). Cortical stroke was modeled as an addition of Gaussian noise to *W*. **c**. Mechanical direction (MD) for the eight muscle groups. **d**. Preferred direction (PD) after optimization with the non-synergy model shown in **a. e**. Cross-validation procedures of non-negative matrix factorization (NNMF) to select the number of muscle synergies. NNMF was applied to the muscle activations of the last 100 trials in the learning simulation with the non-synergy model shown in **a**. Error bars: standard deviation. **f**. Muscle synergies extracted from the non-synergy model. Note that these synergy weights (Sy1–4) were used to define the output of four synergies (*V*) in the synergy model shown in **b**.

#### 2.1.2 Simple synergy model

In the simple synergy model, we added a spinal synergy layer between the cortical and muscle layers (Fig. 1b). Other than the addition of the spinal synergy layer, the synergy model was identical to the non-synergy model. The cortical output to the spinal synergies (*X*) was similarly defined as the non-synergy model (*Z*), but the target number was different. To define the spinal synergies, we extracted muscle synergies from muscle activations produced by the optimized non-synergy model. After optimizing the non-synergy model in the learning simulation of 40,000 trials, we applied a non-negative matrix factorization (NNMF) to the muscle activation patterns during the last 100 trials of the training session. To determine the number of muscle synergies, we performed a four-fold cross validation of NNMF for the different numbers of muscle synergies (1–7), and plotted the averaged variance accounted for (VAF) as a function of the number of muscle synergies (Fig. 1e). VAF was calculated as VAF = 1 – SSE / SST, where SSE is the sum of the squares of residual errors, and SST is the sum of the square differences between each data point and the overall mean. Similar to Hirashima and Oya (2016), we successfully extracted four muscle synergies from their non-synergy network model based on the criterion that the VAF exceeds 0.90. To maintain consistency in the muscle synergy extraction, we set the number of synergies to four for the following NNMF analyses. The synergy weights (Sy1–4) of the non-synergy model were used to define the output of the four spinal synergies (*V*) in the synergy model. In this regard, the synergies explicitly defined in the synergy model were identical to the muscle synergies that were originally extracted from the non-synergy model, which are referred to as original synergies.

#### 2.1.3 Population synergy model

In experiment 2, we further tested another synergy model, in which there were 100 spinal neurons (n = 100). Initially, each group of 25 spinal neurons was provided one of the synergy weights (Sy1–4) for their outputs (*V*). Then, by adding a different proportion of Gaussian noise to the synergy model weights (*V*), we systematically created synergy models with different spinal variation levels. The output weights of each spinal neuron (*V*_*i*_, *i* = 1–100) was obtained by mixing the original synergy weights (Sy_*k*_, *k* = 1–4) with Gaussian noise (*ω*), which was normalized to the norm to be unitary, with a certain proportion (*ρ*):

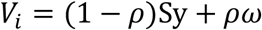

By changing the proportion of noise (*ρ*) from 0 to 1, we systematically investigated the synergy models which had different spinal variations, replicating a variation of the spinal PreM-INs recorded in monkeys performing a precision grip task^24^. The silhouette value was calculated to evaluate the clustering of the spinal neurons with regard to their similarity to the original synergies (Sy1–4)^25^.

### 2.2 Isometric torque production task

We trained each network to perform isometric torque production tasks using a two-joint system (shoulder and elbow)^26^. The target torque combination was chosen from eight possible torque combinations (shoulder flexion [SF], shoulder and elbow flexion [SF+EF], elbow flexion [EF], shoulder extension and elbow flexion [SE+EF], shoulder extension [SE], shoulder and elbow extension [SE+EE], elbow extension [EE], and shoulder flexion and elbow extension [SF+EE]) with the norm of 1 Nm.

### 2.3 Training procedure

The synaptic weight from the input layer to the cortical neurons (*W*) was modified using the error feedback-with-decay algorithm^16,24^:

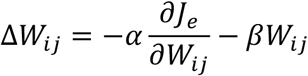

where *α* is the learning rate (*α* = 20) and *J*_*e*_ is the error cost, as calculated by the error vector (*e* = *T* – *τ*) between the output torque and the desired torque: *J*_*e*_ = 1/2*e*^T^*e*. The second term indicates that the changes in synaptic weight due to synaptic memory decay in each step are proportional to the current synaptic weight *W*_*ij*_. The decay rate β was set to 1.0 × 10^−4^, which is much smaller than the learning rate (*α* = 20). By randomly presenting one of the eight target torque combinations in 40,000 trials and modifying the synaptic weight (*W*) after each trial, the network was trained to produce the appropriate output torque.

### 2.4 Stroke model

To simulate the situation of cortical stroke, we added Gaussian noise (∼*N*[0, *σ*^*2*^]) to the cortical synaptic weights (Stroke, Fig. 1a-b and Fig. 5a-b), since previous human imaging studies showed that the long-scale entropy of MEG signal increased in perilesional tissues in stroke patients^27^. We systematically changed the stroke size (σ) from 0 to 10 standard deviations (SD) of the original synaptic weights (*W*). We sampled 200 different stroke cases for each stroke size (σ = 0–10) and evaluated their task performance.

### 2.5 Evaluation of task performance and muscle synergies

To evaluate the stroke effect on task performance and muscle synergies, we used 96 target torque combinations (32 uniformly distributed torque directions × 3 amplitudes [0.5, 1, and 2 Nm]) instead of the eight trained torque combinations. We quantified two types of performance errors, directional error and amplitude error, as a function of stroke size. The directional error was defined as the angle between the target torque vector and the torque vector output by the model. The amplitude error was defined as the difference between the amplitude of the target torque and the model output. Both errors were calculated for each stroke size, and the absolute values were averaged across the 200 stroke cases.

To evaluate the consistency of muscle activation patterns, we quantified the shift in the preferred direction (PD). PD was defined as the direction in which the muscle is maximally active and it was calculated by the summation of the target torque vectors weighted by muscle activity. ΔPD was defined as the absolute shift in PD from no stroke cases (σ = 0).

To evaluate the robustness of muscle synergies in cortical stroke, we quantified two different measures: VAF by the original muscle synergies and similarity of muscle synergies to the original muscle synergies. For these evaluations, we used muscle activations during the same 96 target torque combinations. To evaluate the VAF by the original muscle synergies (Sy1–4), we applied NNMF for muscle activation without updating the synergy weights (Sy1–4). We then calculated the VAF between the observed and reconstructed muscle activity. This measure indicates how much variance in muscle activity can be explained by the original muscle synergies (Sy1–4). We also compared the similarity of muscle synergies extracted in each stroke case with the original muscle synergies. We applied NNMF to muscle activity to extract muscle synergies for each stroke case. For this extraction, we did not use n-fold cross validation, and the number of muscle synergies was fixed at four to allow for a consistent comparison. Then, we calculated the dot product between the extracted synergies and the original synergies. We used the max dot product to determine the similarity of each muscle synergy and averaged the values across the four synergies. This measurement represents the consistency of the muscle synergies between different stroke conditions.

### 2.6 Statistical analyses

To test the significance of the effect of different neural network models on performance measures (directional error, amplitude error, ΔPD, VAF by the original synergies, and similarity with the original synergies), we used the bootstrap method. For the two groups of parameter samples (n = 200), we first calculated the original difference in the mean of two populations (Δmean). Then, we pooled the two populations, resampled two populations of the same size with replacement, and calculated the Δmean. This process was repeated 1,000 times to obtain a baseline distribution of the Δmean. We set the significance limits of this distribution to 0.23 (= 2.5 / number of stroke conditions) and 99.77 (= 100 – 2.5 / number of stroke conditions) percentile that matched a significance level of p < 0.05, with Bonferroni’s correction. All simulations and analytical procedures were performed using MATLAB (MathWorks, RRID: SCR_001622).

## 3. Results

### 3.1 Non-synergy and simple synergy models have comparable training ability

In experiment 1, to test the existence of muscle synergies in the nervous system, we compared the task performance of the neural network model with and without muscle synergies (simple synergy vs. non-synergy models). In the non-synergy model, cortical neurons had random direct connections to the muscles (*Z*). Cortical activation patterns (*W*) were optimized to achieve the model to output the desired shoulder and elbow torques while minimizing the sum of squares of cortical activity (Fig. 1a). As Hirashima and Oya (2016) demonstrated previously, the non-synergy model reproduced a shift in the preferred direction of muscles (Fig. 1d) relative to their mechanical directions (Fig. 1c), and the muscle activation patterns were successfully reconstructed with a linear combination of four muscle synergies (VAF ≥ 0.9, Fig. 1e-f). We refer to the synergies extracted from the non-synergy model as the original synergies (Sy1–4, Fig. 1f). In the simple synergy model, we added four spinal synergies between the cortical and muscle layers (Fig. 1b). We set the output weight of the synergies (*V*) to be the same as the synergy weights extracted from the non-synergy model (Sy1–4, Fig. 1f). Other than the addition of the synergy layer, the simple synergy model was the same as the non-synergy model.

First, we compared the learning performance of the non-synergy and synergy models. Both models were trained to produce eight combinations of shoulder and elbow torques while minimizing the sum of squared cortical activity (i.e., minimizing error cost and motor cost). Fig. 2a shows the trial-dependent changes in the error magnitude averaged across the eight target conditions. After a 40,000-trial training, we found that learning converged in both models, and they reached similar residual errors (Fig. 2c). However, during the training, the learning speeds differed. At the initial phase of the training (Fig. 2a, left), the learning curve was steeper in the synergy model than in the non-synergy model, and the learning rate, fitted by an exponential curve, was higher in the synergy model (Fig. 2b, p < 0.05, bootstrap test, n = 1,000). This indicates that learning progressed faster in the synergy model than in the non-synergy model. We also tested trial-dependent changes in the sum of squared cortical neural activity (Fig. 2d). We found that after training, cortical activity was less for the synergy model than for the non-synergy model (Fig. 2e). Interestingly, despite the prominent difference in cortical activity, the sum of squared muscle activity was the same for both models (Fig. 2f). These results indicate that the synergy model showed faster convergence of learning and smaller neural activity, although both models exhibited similar task performance.

**Fig. 2.**
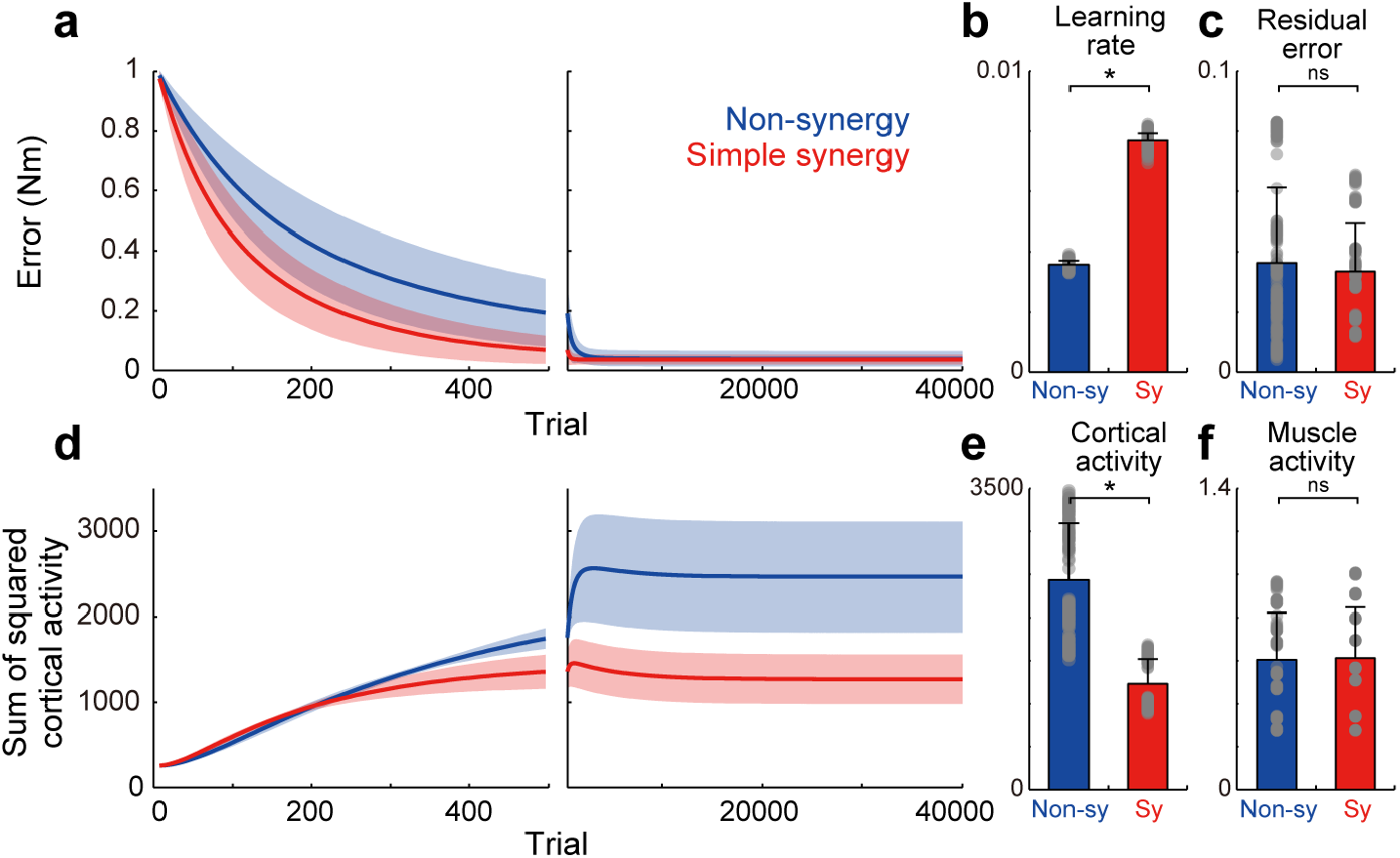
Learning performance of non-synergy and simple synergy models. **a**. Trial-dependent changes in the error magnitude for non-synergy (blue) and simple synergy model (red). Line and shade, mean and standard deviation across the simulations (n=200). Trials of each set of eight targets were averaged for presentation purpose. **b**. Averaged learning speed calculated by fitting of an exponential curve. Gray dots, each simulation. Error bar, standard deviation. **c**. The same format but for residual error at the end of the training. **d**. The same format but for the sum of squared cortical neural activity. **e-f**. The same format but for the sum of squared activity of cortical neurons (**e**) and muscle activity (**f**) at the end of training. *p < 0.05; ns, non-significant, bootstrap test, n = 1,000.

### 3.2 Simple synergy model exhibits a higher robustness against cortical stroke than non-synergy model

We compared the robustness of non-synergy and simple synergy models against cortical stroke. To simulate the situation of cortical stroke, we added random noise to the cortical connections (*W*) of different sizes (σ = 0–10 SD). We tested 200 different stroke cases for each stroke size. Fig. 3 shows three examples of torque output by the non-synergy model (Fig. 3a) and synergy model (Fig. 3b) in no-stroke and in two different sizes of stroke (σ = 1 and 2 SD). Both models showed increasing instability as the stroke size increased. However, systematic differences exist in the distribution of the errors. While the errors of the non-synergy model were distributed almost evenly in all directions (i.e., circular shape), the errors of the simple synergy model were distributed more ovally, and the main axis was along the target direction. For example, for the shoulder flexion target (the rightmost target) with σ = 2SD, the error was distributed horizontally in the simple synergy model (Fig. 3b); however, it was distributed evenly in the non-synergy model (Fig. 3a). To examine the details of the task performance, we evaluated two types of errors: directional error and amplitude error. The directional error is the angle between the target torque vector and the torque vector output by the model, while the amplitude error is the difference between the amplitude of the target torque and the output of the model. Fig. 3c and 3d show an increase in the directional and amplitude errors as a function of stroke size. The figures show that although the amplitude error of both the non-synergy and synergy models increased in a similar manner, the synergy model produced a smaller size of directional error (p < 0.05, bootstrap test, n = 1,000). This result indicates that the synergy model is more robust for cortical stroke to generate torques in the correct direction.

**Fig. 3.**
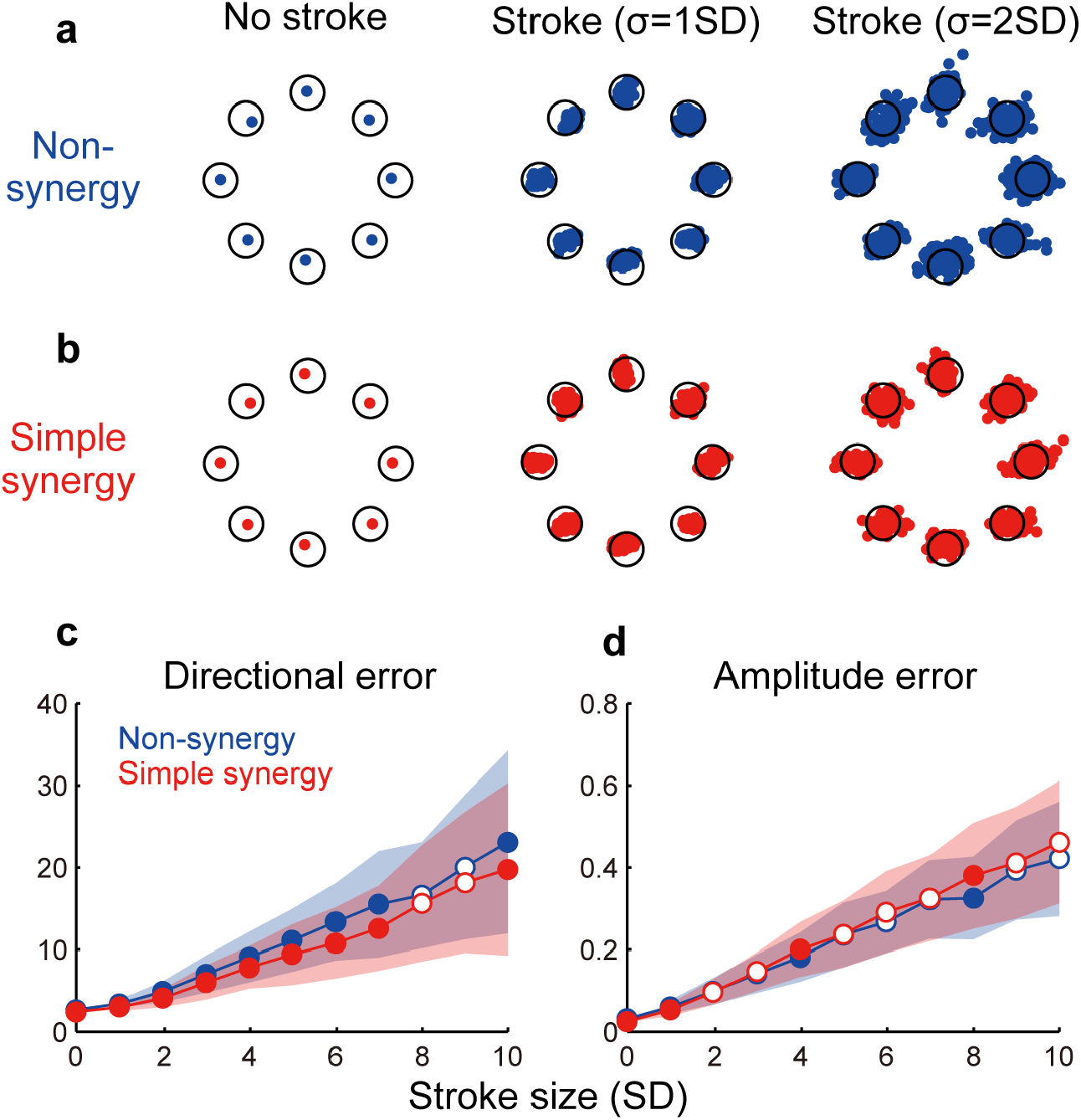
Effect of cortical stroke on task performance. **a-b**. Examples of torque output by non-synergy model (**a**) and simple synergy model (**b**) in no stroke and two different sizes of stroke (σ = 1 and 2 SD). Each dot indicates one stroke case. Only 100 examples are plotted for visualization purpose. Open circle, target torque combination. **c-d**. Size of directional error (**c**) and amplitude error (**d**) as a function of stroke size (σ = 1–10SD). Line and shade, mean and standard deviation across 200 stroke cases. Open and filled circles indicate significant and non-significant differences between two models (p < 0.05, bootstrap test, n = 1,000).

Next, we compared the robustness of muscle activation patterns between non-synergy and simple synergy models. Fig. 4a and 4b illustrate muscle activation for each target direction with non-synergy and synergy models in no-stroke and two different sizes of stroke cases (σ = 3 and 6 SD). While muscle activation of the non-synergy model was severely disturbed by stroke, the synergy model expressed more consistent muscle activation. The shift in PD (ΔPD) was quantified as the absolute difference of the PD from no-stroke cases (σ = 0). Fig. 4c shows that ΔPD was significantly smaller for the simple synergy model than for the non-synergy model (p < 0.05, bootstrap test, n = 1,000). This result indicates that the simple synergy model is more robust for cortical stroke to generate consistent muscle activation.

**Fig. 4.**
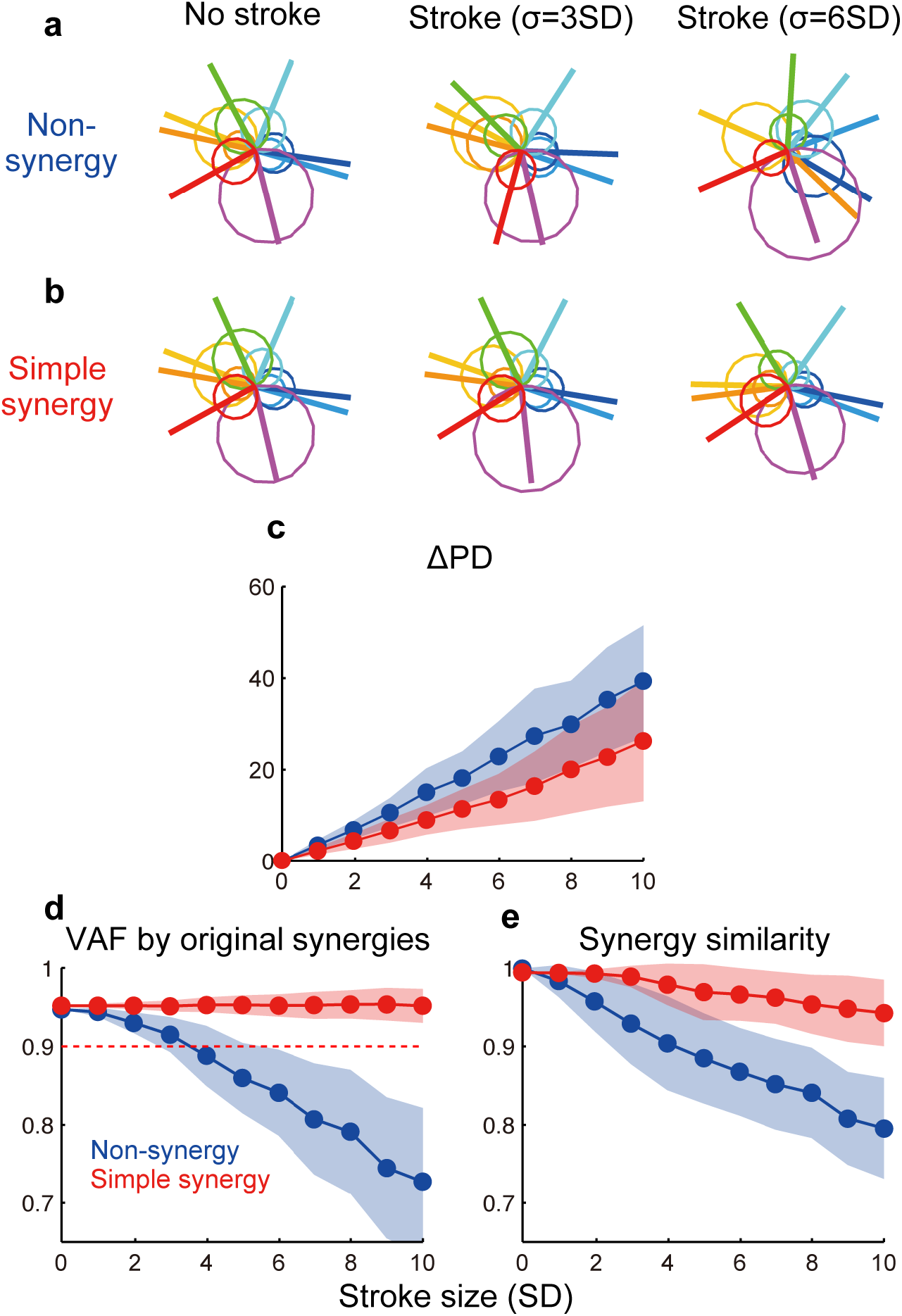
Effect of cortical stroke on muscle synergies. **a-b**. Examples of muscle activation with non-synergy model (**a**) and simple synergy model (**b**) in no-stroke and two different sizes of stroke cases (σ = 3 and 6 SD). Circles indicate muscle activation for each torque direction and lines indicate a PD of muscles. **c**. Averaged shift of PD (ΔPD) from no stroke (σ = 0) condition. **d**. VAF by original muscle synergies extracted from non-synergy model and used for synergy model (Fig. 1f). **e**. Similarity of muscle synergies extracted in each stroke case to the original muscle synergies. The format is the same as Fig. 3c-d.

We further compared the robustness of muscle synergies between non-synergy and simple synergy models. To quantify the robustness of muscle synergies, we compared two measures: 1) how much variance of affected muscle activity can be accounted for by original synergies (VAF by original synergies) and 2) how similar the synergies extracted from the affected muscle activity was to the original synergies (similarity with original synergies). Fig. 4d and 4e demonstrate that while both measures steeply decrease in the non-synergy model, they remain high in the simple synergy model (≥0.90, Fig. 4d, red dotted line), even when the stroke size increased to 10 SD. The difference in the effect of stroke size on each model was significant for both measures (p < 0.05, bootstrap test, n = 1,000). These results indicate that the synergy model more robustly generates consistent muscle synergies under stroke conditions.

### 3.3 Population synergy model with a moderate spinal variation exhibits a comparable robustness to simple synergy model

We examined how the synergies could be organized in the spinal layer. The simplest model is that in which each synergy is represented by a single neuron or unitary “module” (simple synergy model, Fig. 1b and 5a). However, our previous physiological study showed that a muscle field of individual spinal PreM-INs did not completely match muscle synergies but showed substantial variation^13^. Therefore, a more plausible model can be developed in such a way that a synergy is represented by a population of neurons with some variation (population synergy model, Fig. 5b). To test this scenario, in experiment 2, we compared the robustness of these synergy models against the introduction of cortical stroke. First, we systematically changed the size of the variation of synergies by changing the proportion (*ρ*) of Gaussian noise to the synergy weights (*V*). We found that spatial clustering of the output weights of each spinal neuron based on the similarity to the original muscle synergies decreased as the variation level *ρ* increased (Fig. 5c). The level of clustering was evaluated using silhouette values (0.93, 0.81, and 0.56 for *ρ* = 0.2, 0.3, and 0.4, respectively)^25^. Importantly, our previous electrophysiological study demonstrated that the similarity of output projection of spinal PreM-INs to extracted muscle synergies showed a spatial clustering and their silhouette values ranged from 0.79 to 0.87 (Fig. 2b of Takei et al. 2017^13^). This silhouette value was comparable to that of the population synergy model, with *ρ* = 0.3.

**Fig. 5.**
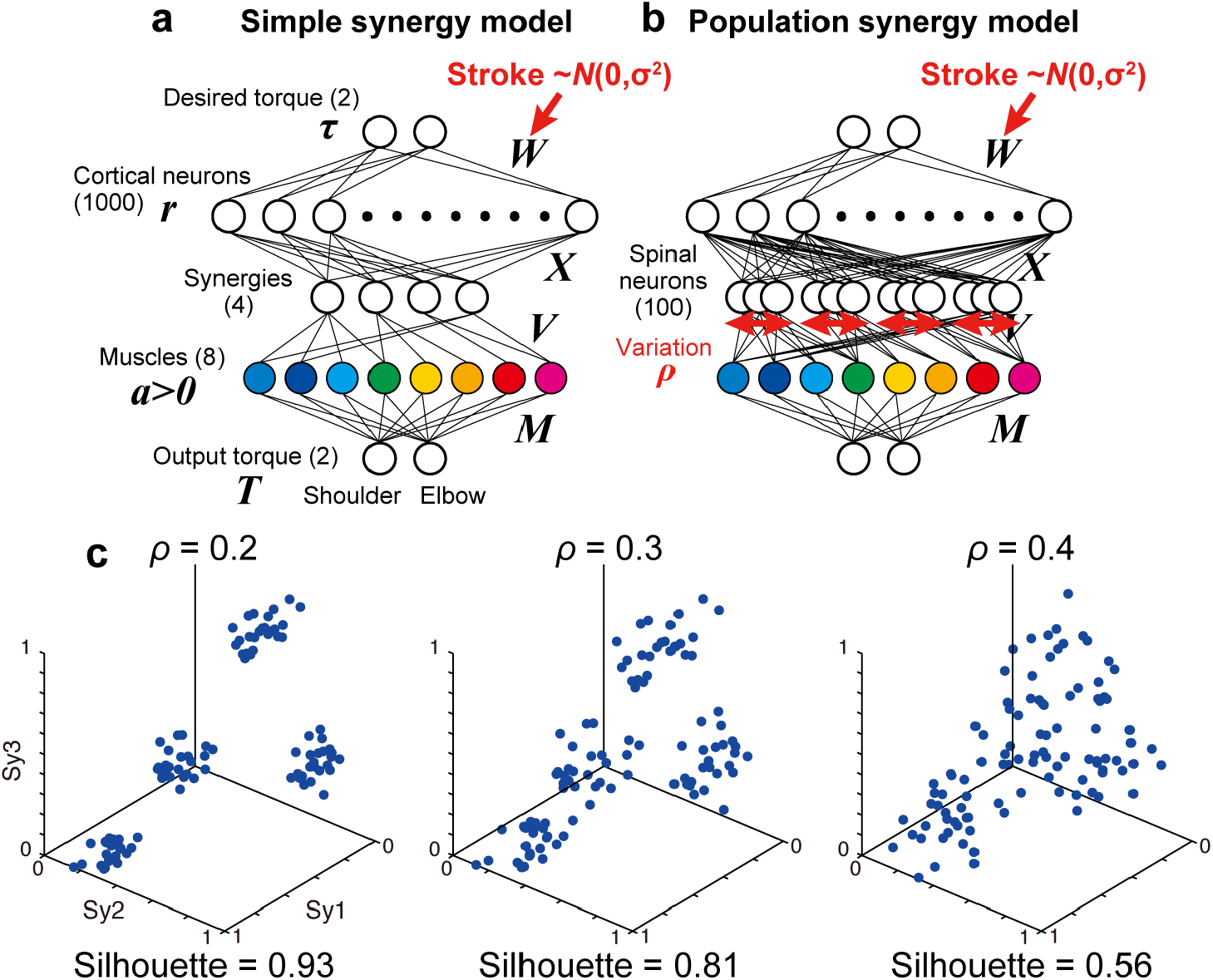
Simple and population models for muscle synergies. **a-b**. Neural network models with simple synergies (**a**, same as Fig. 1b) and population synergies with some variations (**b**). **c**. Spatial similarity of output weights of each synergy (*V*) with the original muscle synergies with different spinal variation levels (*ρ* = 0.2, 0.3, 0.4).

Then, we compared the robustness of task performance and muscle synergies in cortical stroke between the simple and population synergy models. Fig. 6a-e shows the change in task performance, ΔPD, and muscle synergies as an effect of cortical stroke. As a reference, we also plot the results of the non-synergy model (Fig. 6a-e, gray rectangles). In general, results of the population synergy models spanned from the results of the simple synergy model to those of the non-synergy model. For example, the directional error of the population synergy model was similar to the simple synergy model when the spinal variation was smaller (*ρ* = 0.0–0.2, Fig. 6a-e, red lines), while it was similar to the non-synergy model when the variation was larger (*ρ* = 0.8–1.0, blue lines). This gradual change was observed for directional error, ΔPD, and two muscle synergy measures (p < 0.05, bootstrap test, n = 1,000), but not for the amplitude error (Fig. 6b). Critically, the robustness of the population synergy models remained almost at the same level as the simple synergy model until the spinal variation level was increased to 0.3 (Fig. 6c-e, red-yellow), and then these measures steeply decreased when the variation increased further (green-blue). These results indicate that the population synergy model with moderate variation exhibited robustness against cortical stroke comparable to that of the simple muscle synergy model. These results suggest that the muscle synergies are not necessarily implemented as unitary “modules” as they were often modeled before, but they can be implemented as the population of spinal neurons with moderate variation.

**Fig. 6.**
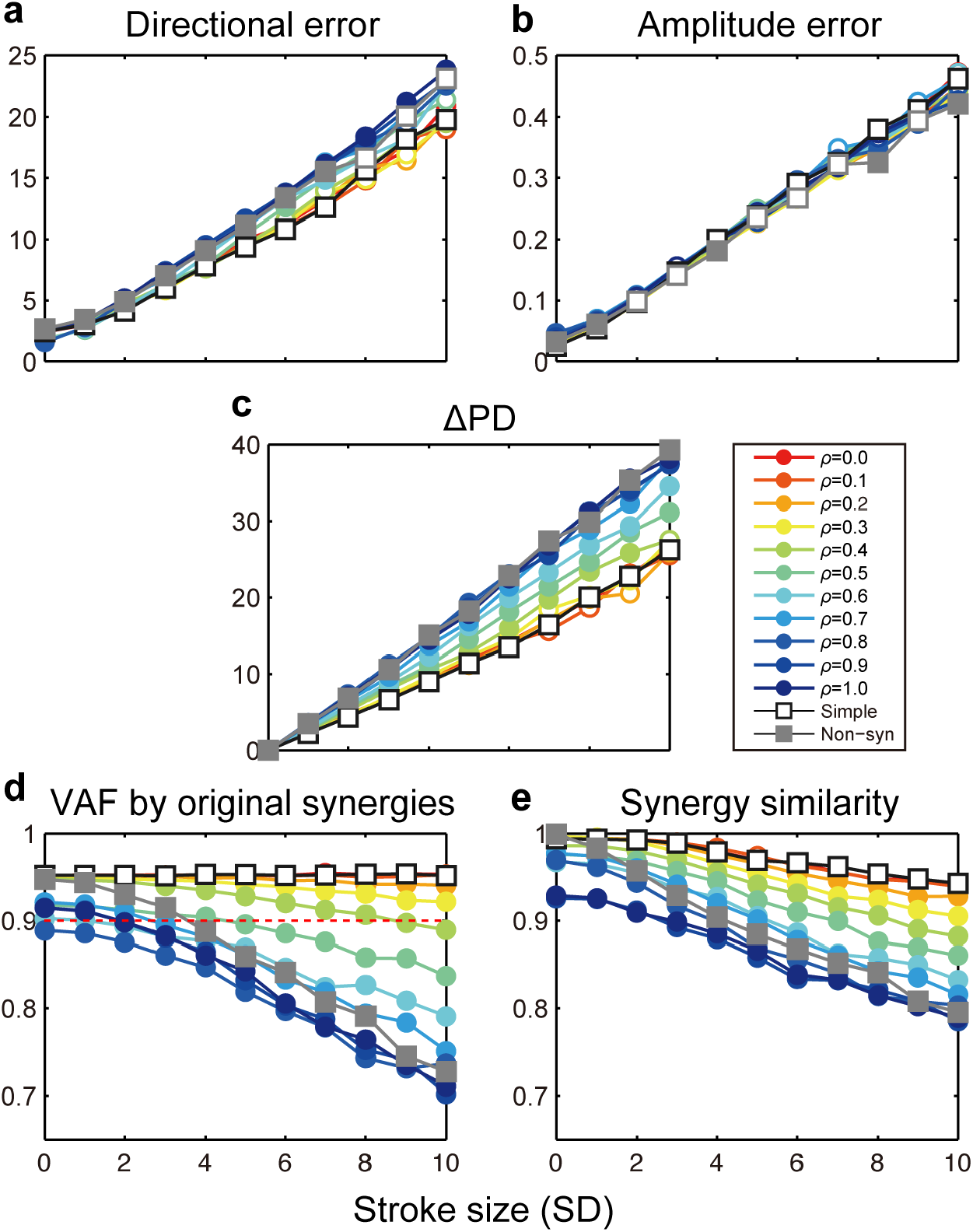
Stroke effects on task performance and muscle synergies with simple and population synergy models. Change in directional error (**a**), amplitude error (**b**), shift of PD (**c**), VAF by the original synergies (**d**), and similarity with the original synergies (**e**) as a function of the size of cortical stroke (SD). Open and filled circles indicate a significant and non-significant difference from the simple synergy model (open square, p < 0.05, bootstrap test, n = 1,000).

## 4. Discussion

In this study, we compared different neural network models with a focus on their robustness to cortical stroke. In experiment 1, we compared two types of neural network models: one with random connections (non-synergy model) and the other with a smaller number of spinal synergies (simple synergy model). After optimization, we found that both models achieved comparable task performance and similar muscle synergies, confirming the prediction of a previous study^16^. Interestingly, despite the similar task performance after the training, the learning rate was higher and cortical activity was smaller for the synergy model (Fig. 2a-f). This indicates that although the output performances were similar, the synergy model achieved more efficient muscle control than the non-synergy model.

Then, we tested the robustness of the model performance against cortical stroke, which was modeled as the addition of noise to the cortical layer. The results demonstrated that the synergy model exhibited 1) smaller directional error in torque production and 2) higher consistency of muscle synergies with the original muscle synergies. This robustness of muscle synergies was consistent with observations in human stroke patients^22,23^. Cheung et al. (2009) compared muscle synergies extracted from the unaffected and affected arms of stroke patients with moderate-to-severe unilateral ischemic lesions in the frontal motor areas^22^. Their results demonstrated that most of the patients showed muscle synergies that were strikingly similar between the unaffected and affected arms despite differences in motor performance between the arms. The similarity evaluated by a dot product between muscle synergies from the arms ranged from 0.80 to 0.95 with an average of 0.90. Our results showed that the similarity of muscle synergies with the original synergies was maintained at >0.90, even with a severe stroke in the simple synergy model (Fig. 4d, red dotted line). The consistency of our results with previous findings suggests that the existence of a synergy layer is an essential factor in explaining the robustness of muscle synergies in stroke conditions.

We further explored a biologically plausible model for the neural implementation of muscle synergies. Our previous study demonstrated that spinal PreM-INs have divergent output projections to multiple hand and arm motoneurons^20,21^ and their spatial distribution corresponds to the spatial weight of muscle synergies^13^. However, although the output projection of PreM-INs corresponded to muscle synergies as a population, there was a substantial variation in the projection patterns across individual PreM-INs. From these observations, we hypothesized that each muscle synergy is not represented by a unitary “module,” but by a population of PreM-INs^13^. Consistent with this hypothesis, the present results demonstrated that the population synergy models with a spinal variation exhibited comparable robustness of muscle synergies with the simple synergy model against cortical stroke^22,23^. The similarity of muscle synergy to the original synergies remained high (>0.90, Fig. 6d, red dotted line) even when the spinal variation level was increased up to *ρ* ≤ 0.3 (Fig. 6d-e, red-yellow). Importantly, the variation of spinal PreM-INs reported in our previous study^13^ corresponded to *ρ* = 0.3, according to the silhouette value calculated from the cluster analysis for spinal PreM-INs (Fig. 2b of Takei et al. 2017^13^). This variation level was within the range where the present models preserved the robustness of muscle synergies against cortical stroke. These findings suggest that the population of spinal neurons with moderate variation is a biologically plausible model for the neural implementation of muscle synergies.

This finding has several implications for the low-dimensional control of the limb and muscles. First, muscle synergies do not necessarily reduce the degrees of freedom (DOFs) of motor control because in the population synergy model, the DOFs of motor control in the spinal layer (i.e., a number of spinal neurons, n = 100) are not smaller than the number of muscles (n = 8). Previously, muscle synergy has often been modeled as a linear combination of a smaller number of modules and interpreted as a simplified strategy for motor control. However, the spinal PreM-IN showed substantial variation between neurons to contradict such a simple view of modular control. The present simulation results suggest that the populational representation of muscle synergies in spinal neurons does not reduce the DOFs but increases the robustness of muscle coordination in a task space. A similar task-control space has been identified as neural manifolds in the population activity of motor cortices^28-30^. It is interesting to investigate how the cortical manifold and spinal synergy space interact and are hierarchically organized in the motor system.

Our model of spinal population with a moderate variation in muscle synergy also provides implications for motor learning and development of motor coordination. Previous studies have demonstrated that motor tasks compatible with the original muscle synergy are learned faster than tasks incompatible with muscle synergies^31^. Interestingly, the results also showed that although the learning rate was slow during the incompatible task, learning progressed to explore new synergies. During this exploration process, the variation in spinal neurons may be involved in changing the muscle coordination patterns. It is also interesting to understand how muscle synergies are obtained during development. In the sensory system, it is known that the synaptic connection in primary sensory cortices increases after birth till it reaches a maximum, then prunes and decreases as neural selectivity increases^32^. This pruning may also occur during the development of the motor system. How the variation of spinal and motor systems changes during development to optimize the low-dimensional control space in the spinal layers should be investigated in future studies.

In conclusion, our network simulation showed that muscle synergies could be implemented as a population of PreM-INs with a moderate variation rather than unitary “modules.” This view provides new insight into understanding the mechanism and functional relevance of how the CNS controls the redundant motor system.

## Author contributions

T.T. designed the study. Y.S. and T.T. carried out the experiments. Y.S. and T.T. analyzed the data. Y.S., M.H. and T.T. wrote the manuscript.

## Conflict of interest

The authors declare no conflict of interest.

## Acknowledgements

This work was supported by Grants-in-Aid (19H03975, 19H05311, 21H00309) from the Ministry of Education, Culture, Sports, Science and Technology of Japan (MEXT), The Fusion Oriented REsearch for disruptive Science and Technology (FOREST) program of Japan Science and Technology Agency (JST), The Uehara Memorial Foundation, The Naito Foundation, and The Takeda Science Foundation (T.T.).

## Notes

### Competing Interest Statement

The authors have declared no competing interest.

